# Engineered balanced lethal systems for partial suppression or enhancement of wild populations

**DOI:** 10.64898/2026.06.29.735349

**Authors:** Katie Willis, Austin Burt

## Abstract

Genetic interventions to modify wild population densities are typically framed around pest suppression, with parallel approaches for boosting beneficial or endangered populations remaining largely undeveloped. Imposing a sustained but non-eliminative genetic load could in principle address both objectives, but existing designs rely on genes with intermediate fitness effects whose loads are difficult to predict under field conditions. Here we describe engineered balanced lethal systems, in which CRISPR-based gene drive establishes two complementing recessive-lethal alleles at a single locus, producing a sustained 50% load through Mendelian segregation. Modelling shows these systems spread from small releases, and that the resulting population-level consequences depend on density regulation and on the timing of lethality: the same 50% load can suppress pests, boost populations of beneficial or endangered species, dampen boom-bust cycles, or raise effective population size. Additional systems at independent loci scale the effect in stepwise increments, and a split-drive variant localises it geographically. These results demonstrate that gene drives imposing genetic load can be expanded beyond elimination, to support and preserve beneficial and endangered populations.

## 1. INTRODUCTION

Modifying the density of wild populations is a shared goal across a range of contexts: to suppress the numbers of harmful populations like disease vectors, agricultural pests or invasive species; or to increase the numbers of beneficial species such as pollinators, biocontrol agents or those at risk of extinction. Genetic interventions, in which modified individuals of the same species are released into wild populations, are one approach. The most widely used example is the sterile insect technique, in which individuals sterilised by radiation, precise genetic modification or cytoplasmic incompatibility mate with wild females and reduce population reproductive output (Alphey et al., 2010, Dyck et al., 2021). More recently, CRISPR-based gene drives have been developed that bias their own inheritance above the 50% Mendelian expectation, allowing them to increase in frequency from rare over multiple generations (Raban et al., 2023, Grilli et al., 2021, Hay et al., 2021). This biased transmission can carry linked alleles that reduce survival or fecundity to high frequency despite their negative fitness consequences, imposing a genetic load that changes population size and dynamics. Because they can spread from rare, drive alleles can have large and sustained effects on populations from small releases, potentially across a species’ range wherever there is gene flow.

The two main molecular approaches to engineering gene drives have been homing-based systems and toxin-antidote systems, both typically implemented using CRISPR-based technologies (Burt, 2003, Champer et al., 2020b, Oberhofer et al., 2019). A range of drive designs have been proposed that vary in how they spread: some are self-sustaining from rare (Burt, 2003, Champer et al., 2020b, Oberhofer et al., 2019); some spread from rare but can be confined to target populations through sequence-specific targeting (Sudweeks et al., 2019, Willis and Burt, 2021, Geci et al., 2022); some require release above a threshold frequency to spread (Champer et al., 2020a, Davis et al., 2001, Akbari et al., 2013); and some drive only temporarily (Noble et al., 2019, Oberhofer et al., 2021). In addition, a range of non-driving designs have been proposed that do not increase from low frequency and are either selectively neutral (Burt and Deredec, 2018, Willis and Burt, 2025a) or expected to disappear from the population (Thomas et al., 2000, Kandul et al., 2019, Alphey, 2002, Willis and Burt, 2025b). Despite this breadth of mechanism, designs intended to change population size have focused almost exclusively on population suppression, and on maximising impact to achieve elimination of the target population.

Complete elimination of the target population, however, is not always the desired outcome. Some pest species are harmful only at high density but are benign or even beneficial at lower numbers, and populations with cyclical dynamics may be problematic only during periodic booms. For such cases, a sustained and predictable partial genetic load may be a more appropriate intervention. Existing routes to partial load are limited. Load-inducing alleles have typically been generated by disrupting essential genes whose knockout results in lethality or sterility (Kyrou et al., 2018, Hammond et al., 2016, Fuchs et al., 2021, Verkuijl et al., 2025). The most efficient of these designs combine such knockouts with self-sustained gene drive, but these aim for maximum genetic load and population elimination. In principle, partial genetic load could be achieved by targeting genes where knockout has intermediate rather than lethal or sterile fitness consequences (Burt, 2003, Deredec et al., 2008), but this raises the problem of predicting the fitness consequences of knocking out a non-essential gene under field conditions. Alternatively, partial load could be imposed using designs that drive only temporarily, or those that do not drive at all and instead persist at the release frequency or decline, but these designs typically require large or repeated releases, making them less efficient than self-sustaining drive-based approaches.

More broadly, the population-level consequence of a given genetic load depends on how density is regulated in the target population and on when in the life cycle the load is imposed. For species where density-dependent regulation occurs at the juvenile stage, a genetic load imposed after the juvenile stage will suppress the population more than one imposed before (Alphey and Bonsall, 2014, Deredec et al., 2011). Indeed, Alphey & Bonsall (2014) have shown that in species where over-compensatory density dependence occurs (where a reduction in the number of juveniles can lead to an increase in the number of surviving adults (Hassell et al., 1976)), a small genetic load before density dependence can actually increase the number of adults and dampen the cyclical population dynamics that can arise with over-compensation. In principle, the same genetic load that suppresses one population could stabilise fluctuations or even increase the size of another, depending on the underlying density regulation. This broader range of outcomes has received less attention than suppression alone.

Here we propose a strategy that imposes a sustained, predictable partial genetic load by using gene drives to establish a balanced lethal system. Balanced lethal systems are naturally occurring genetic architectures in which two recessive lethal alleles at a single locus each compensate for the function lacking in the other (Wielstra, 2020): heterozygotes carrying both alleles are fully viable, while homozygotes for either allele are lethal. In a population fixed for the two alleles at equal frequency, all adults are heterozygous; random mating produces 50% homozygous offspring that do not survive and the population sustains a consistent 50% genetic load. Despite this reproductive wastage, balanced lethal systems have become fixed in natural populations including *Triturus* newts, some insects (*Drosophila tropicalis* and *Tribolium castaneum*) and some plants (Wielstra, 2020). However, due to their fitness costs, they are not expected to spread from rare without an additional mechanism to bias their inheritance. Using mathematical modelling, we show how CRISPR-based homing drives can establish a balanced lethal system in a target population, and we characterise its consequences for population size and dynamics. We demonstrate that, depending on the density regulation of the target population and the time at which the balanced lethal is expressed, a sustained 50% genetic load can suppress populations, dampen oscillatory or chaotic fluctuations in population size, or increase the equilibrium population size and effective population size (Ne). These results open the possibility of applications in pest control, conservation of threatened populations, and management of species that are harmful only during periodic booms. Stepwise intervention via independent loci allows the intensity of load to be tuned, and a split-drive variant permits localised deployment without range-wide spread. Because the load our designs impose arises from Mendelian segregation between two well-characterised lethal knockouts, rather than from the field fitness of a disrupted non-essential gene, it is more predictable than approaches relying on genes with intermediate or context-dependent effects.

## 2. RESULTS

### Establishing a balanced lethal system using homing gene drives

We first consider how a balanced lethal system could be engineered by leveraging intragenic complementation, whereby two mutant alleles of the same gene, each disrupting a different essential function, together restore wild-type function *in trans* (Fincham, 1968). Intragenic complementation can arise through several mechanisms, including proteins with distinct functional domains (e.g. *rudimentary* in *Drosophila*; Carlson (1971)) and genes producing distinct isoforms through alternative splicing (e.g. *Sex-lethal* in *Drosophila*; Salz et al. (1987)). We illustrate our approach with one specific architecture based on alternative splicing: two autonomous homing constructs inserted into separate exons of the same essential autosomal gene, where each exon is specific to a different essential protein isoform generated through alternative splicing (Figure 1a). Each construct contains a Cas9 and a gRNA that targets the wild-type sequence at its own insertion site, enabling homing via homology directed repair. The insertion of each construct eliminates one isoform while preserving the other, so that homozygotes for either construct lack one essential isoform and are lethal, while trans-heterozygotes are viable through complementation of the two isoforms, one from each chromosome (Figure 1a). Each construct also carries a recoded version of the target sequence at the other exon, making it resistant to cleavage and preventing homing between the two constructs in trans-heterozygotes, which would otherwise generate haplotypes lacking both isoforms. The two target exons must be sufficiently close that the recoded site falls within the gene conversion tract during homing, ensuring it is co-transferred with the construct.

**Figure 1.**
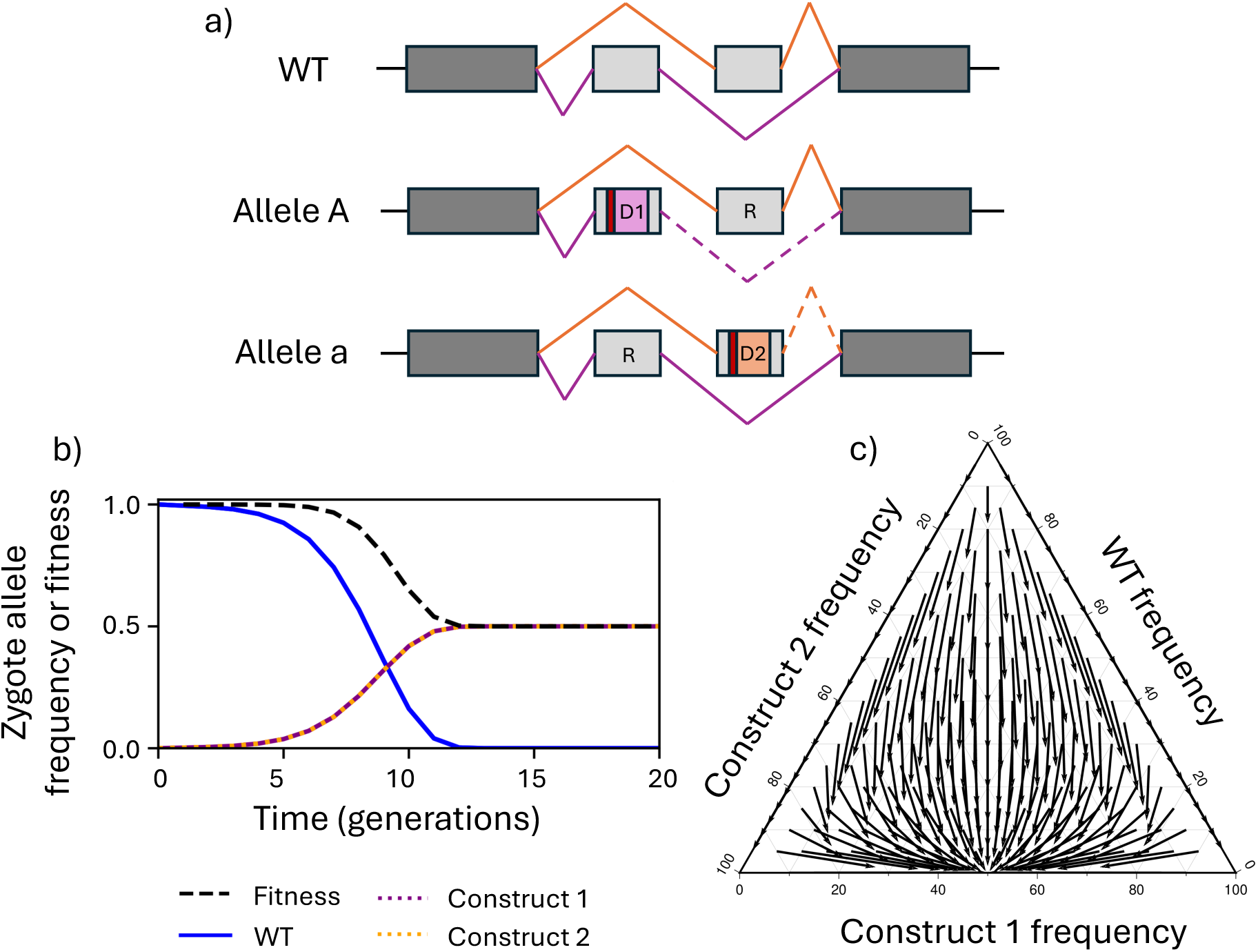
**a)** A schematic of the wild-type allele (+) and two drive alleles (A and a). Boxes represent shared (dark grey) and mutually exclusive (light grey) exons specific to different isoforms. Orange and purple lines indicate alternative splice patterns used to generate two different essential isoforms. Homing constructs (D1 and D2) are inserted into exons specific to different isoforms, disrupting expression of one isoform while preserving the other. Dashed lines indicate disrupted splice patterns due to upstream D1 and D2 insertions. Homozygotes for either allele (A/A or a/a) are lethal since they lack one of the two essential isoforms, whilst heterozygotes (A/a) survive since each isoform is expressed from one chromosome. Each homing allele also includes a recoded target site (R) for the intact exon, making it resistant to cleavage and preventing homing in trans-heterozygotes. **b)** Simulation of the establishment of a balanced lethal system using two homing gene drives released in heterozygous males carrying one copy of each type at generation zero at 1% of the initial male population size. Homing constructs are both inserted into the same locus and designed to home into the wild-type version of that locus. Each construct causes lethality when homozygous but has no fitness effect when paired with the wild-type or the opposing construct. Lines show allele frequencies of the wild-type (blue), construct 1 (purple) and construct 2 (orange) over time. **c)** Change in allele frequency in a single generation for a range of starting combinations of the wild-type and two homing constructs. From all starting points that contain at least some proportion of both constructs, the system tends to a 50:50 ratio of both constructs and eliminates the wild-type. All simulations assume 100% homing efficiency and the absence of migration.

Using a deterministic population genetic model, we simulate the release of trans-heterozygous males into an isolated population, assuming complete homing efficiency and that homozygosity for either construct results in lethality. Following a 1% release of trans-heterozygous males, both constructs spread from rare due to homing and reach an equilibrium allele frequency of 50% each (Figure 1b, purple and orange). The wild-type allele is eliminated by homing, and since homozygotes for either construct are lethal, the adult population consists entirely of trans-heterozygotes. Each generation, half of all offspring are homozygous for one or the other construct and do not survive to adulthood, so the population sustains a consistent 50% genetic load. This equilibrium is robust to variation in initial conditions: even if the constructs are released at unequal frequencies, or if allele frequencies are perturbed by stochastic drift or short-term immigration of wild-type individuals, the system will converge on the same 50:50 equilibrium of the two constructs, eliminate the wild-type, and establish a sustained 50% load, provided both the constructs remain present in the population (Figure 1c and Supp. Figure 1a).

Because the balanced lethal system depends on the knockout phenotype of each allele rather than on the ability to home, it is not necessary for both constructs to autonomously home. The second allele can simply be a non-driving knockout of its respective exon, carrying a target-site modification that prevents its own conversion by homing. In this design, the homing construct spreads from rare through biased inheritance, progressively eliminating the wild-type allele at its target exon. As it does so, it generates indirect selection favouring the complementary knockout: individuals carrying both alleles are viable trans-heterozygotes, while those homozygous for the homing construct are lethal. The non-driving allele therefore increases in frequency, lagging behind the homing construct, until both alleles reach a 50:50 equilibrium and the balanced lethal system is established (Supp. Figure 2).

### Impact on population size and dynamics

The consequence of a sustained 50% load depends on how the target population is density-regulated and on when in the life cycle the construct’s fitness cost acts. Depending on these two factors, the same 50% load can suppress the population, increase it, or dampen its fluctuations without changing its mean size. We modelled the impact of releases using a simple population and genetics model with four life stages: adults mate to produce zygotes, zygotes survive to hatchlings, hatchlings undergo density dependent competition following a generalized Beverton Holt model (Beverton and Holt, 2004, Maynard Smith and Slatkin, 1973, Bellows, 1981, Alphey and Bonsall, 2014), and survivors pupate, before emerging as adults. We varied two parameters: Rm, the intrinsic rate of increase (how fast the population grows at low density), and B, the strength of density dependence (how sharply mortality increases with density).

When the constructs’ fitness cost acts after density-dependent mortality (Figure 2, purple), a 50% load always suppresses the population, with greater suppression at lower Rm or lower B. When the cost acts before density-dependent mortality (Figure 2, green), the same 50% load can either suppress or increase the population, depending on B. At B = 1, mortality scales linearly with density, and the load is suppressive under both timings. At B > 1 (overcompensation), small changes in density produce disproportionate changes in survival, and removing individuals before density dependence acts relaxes competition enough that more of the remaining juveniles survive to adulthood than would have done otherwise, and the population grows. The timing of lethality relative to density-dependent mortality therefore determines the ecological outcome, as has been shown for other gene drive strategies (Alphey and Bonsall, 2014): under the same density regulation, different timings can produce opposite effects on population size (e.g. Figure 2, Rm = 6, B = 2.5).

**Figure 2.**
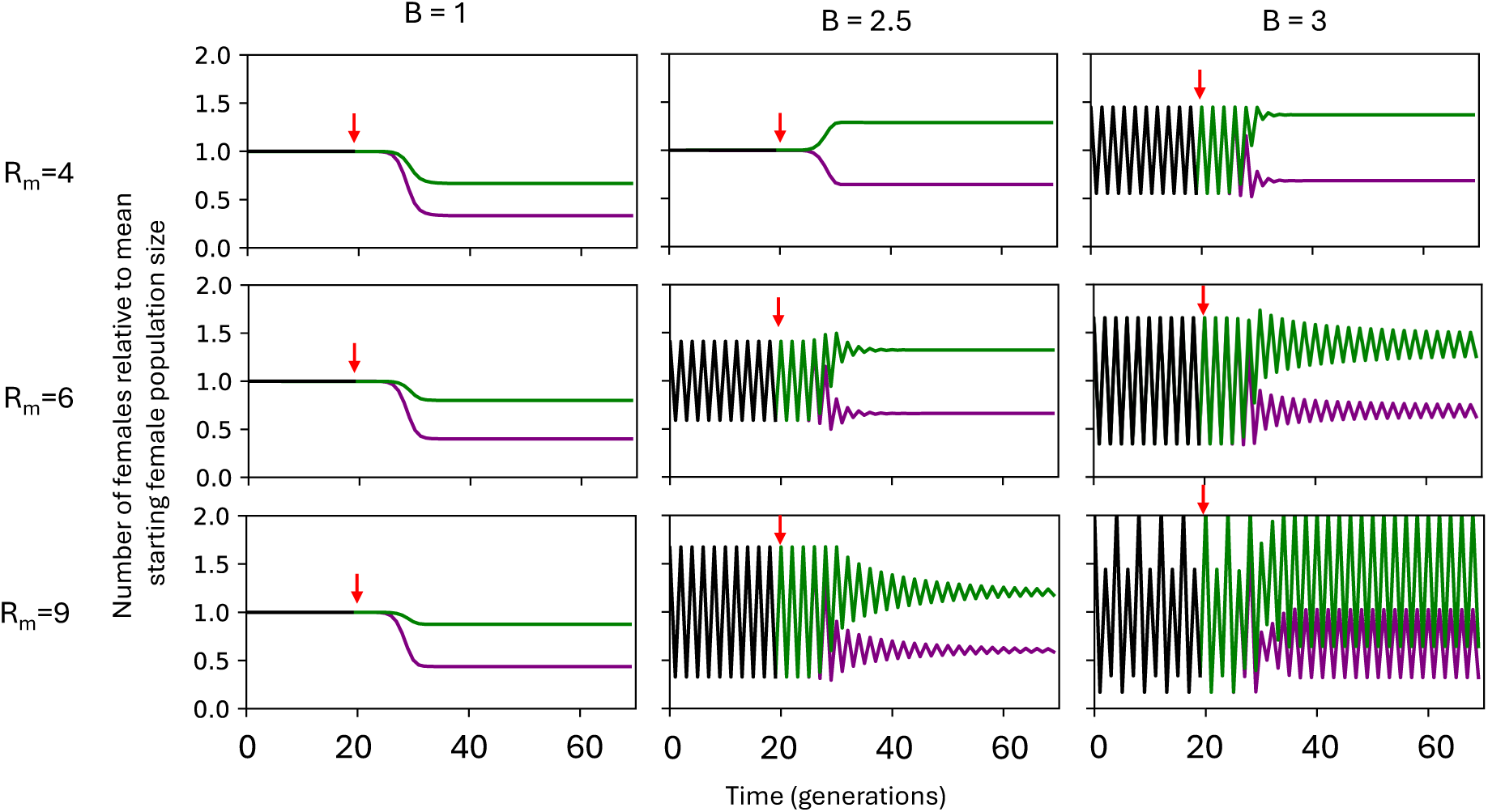
Impact of a balanced lethal system on population size and dynamics. Each panel shows the number of females relative to the arithmetic mean pre-release female population size over time (generations), for populations varying in intrinsic rate of increase (Rm = 4, 6, 9; rows) and strength of density dependence (B = 1, 2.5, 3; columns). Red arrows indicate the release of trans-heterozygous males at 1% of the initial male population size. Black lines show pre-release population dynamics. Green lines show population dynamics when construct-induced lethality occurs before density-dependent mortality. Purple lines show population dynamics when lethality occurs after density-dependent mortality. When B = 1, a 50% load results in suppression under both timing scenarios, with greater suppression at lower Rm. When B > 1 (overcompensation), the timing of lethality determines whether the population is suppressed (purple) or increased (green), and oscillatory dynamics are dampened under both scenarios. All simulations assume 100% homing efficiency, idealised fitness effects and an isolated population.

Some populations exhibit oscillatory or chaotic dynamics when density regulation is strongly overcompensating, even in the absence of seasonal influences. In these cases, inducing a 50% load dampens the fluctuations under both lethality timings, whether the overall effect on mean population size is to increase or decrease it. This can reduce peak population size even when the arithmetic mean of the population increases (e.g. Figure 2, Rm = 9, B = 2.5) and raise the minimum population size regardless of the direction of change in the mean, reducing susceptibility to stochastic extinction. The effective population size (Ne) at loci unlinked from the balanced lethal system, which is set by the harmonic rather than the arithmetic mean of census size, can therefore increase even when the arithmetic mean census size is reduced, increasing genetic diversity and reducing inbreeding depression (Supp. Figure 3).

If a single sustained 50% load is insufficient for the desired outcome, additional balanced lethal systems can be established at independent loci, each imposing a further 50% load and allowing for stepwise changes in population dynamics. Sequential releases at two unlinked loci produce incremental suppression, enhancement, or stabilisation of the population, matching the direction of the first release, with the second producing a comparable additional shift (Figure 3). This demonstrates that the approach is additive and can be tuned by the number of systems established.

**Figure 3.**
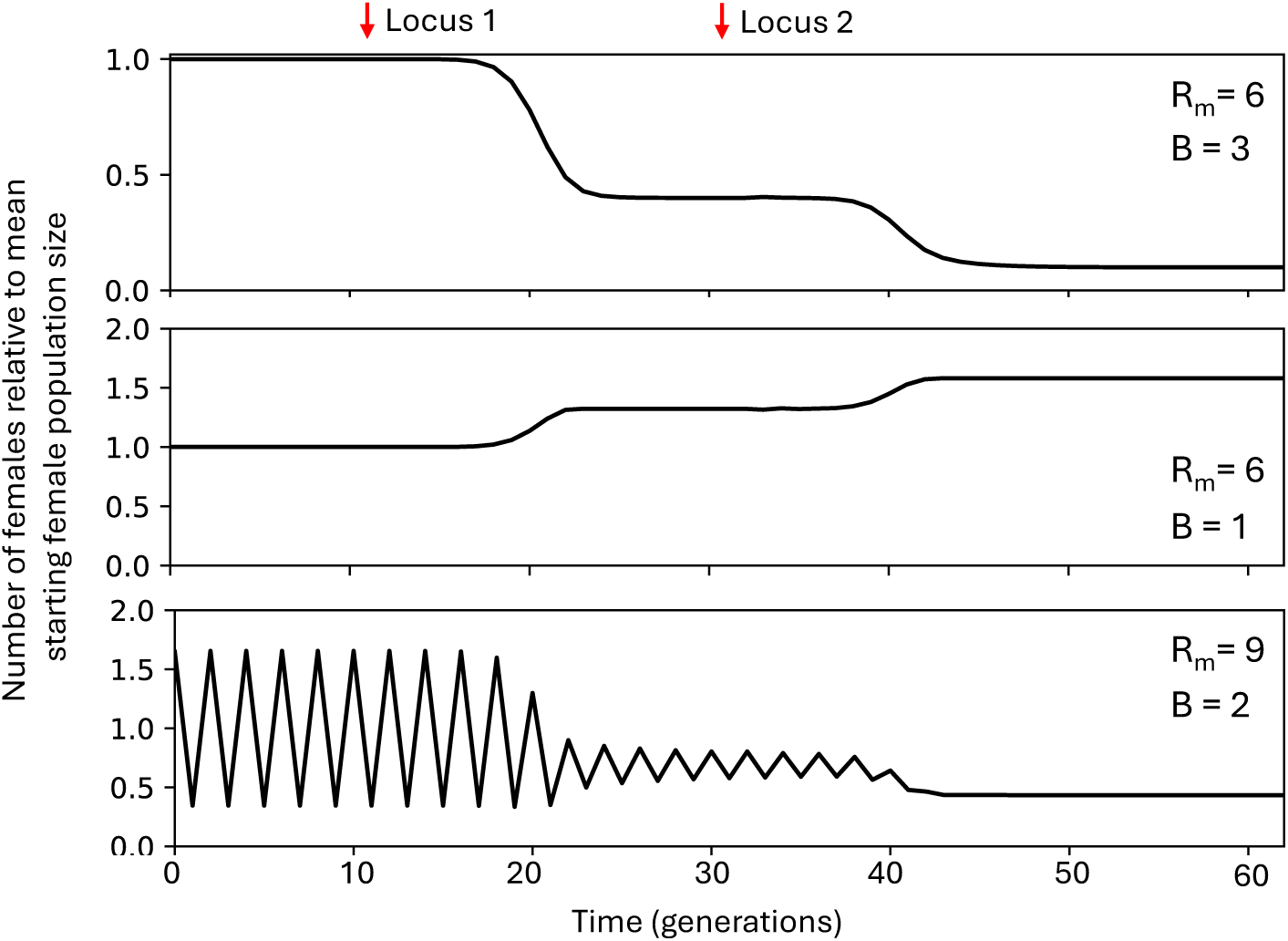
Stepwise changes in population dynamics through sequential establishment of balanced lethal systems at independent loci. Each panel shows the number of females relative to the mean pre-release female population size over time (generations). Red arrows indicate the release of trans-heterozygous males, first for Locus 1 and then for Locus 2. Top panel: stepwise suppression, where each balanced lethal system further reduces population size (Rm = 6, B = 3, lethality acting after density dependent mortality). Middle panel: stepwise enhancement, where each system further increases population size (Rm = 6, B = 1, lethality acting after density dependent mortality). Bottom panel: an oscillating population in which the first balanced lethal system dampens fluctuations and the second further stabilises and reduces the population (Rm = 9, B = 2, lethality acting before density dependent mortality).

The same 50% load can thus suppress pest or vector populations, boost populations of beneficial species or those at risk of extinction, reduce peak densities of species that are damaging only during boom generations, or increase effective population size in species where genetic diversity is limiting, with the outcome depending on the density regulation of the target population and the timing at which fitness costs are imposed.

### Stability

The results above assume idealised conditions: an isolated population and intact genetic constructs maintaining function indefinitely. We now examine how wild-type immigration and homing-associated loss-of-function affect the equilibrium load, and how release conditions shape the transient dynamics during establishment.

A constant influx of wild-type individuals from elsewhere reduces the equilibrium genetic load, because immigration maintains wild-type alleles in the population and limits the frequency the constructs can reach. The system is nonetheless robust to low levels of immigration, with substantial load maintained at moderate immigration rates (Figure 4a). Each genetic component (gRNA or Cas9) of the constructs may also degrade in function over time, reducing or eliminating their ability to home. In the absence of immigration, this has no impact on the equilibrium load, provided the degenerate alleles maintain their fitness effects in homozygotes and complementarity in heterozygotes (Figure 4a). This is because the balanced lethal system depends on the knockout phenotype of each allele, not on the drive machinery: once wild-type alleles have been eliminated, further homing is no longer required and constructs that have lost drive function continue to contribute to the balanced lethal system alongside the intact ones. When immigration and loss of function occur together, the impact on equilibrium load can exceed that of immigration alone, because constructs that no longer home cannot eliminate incoming wild-type alleles (Figure 4a).

**Figure 4.**
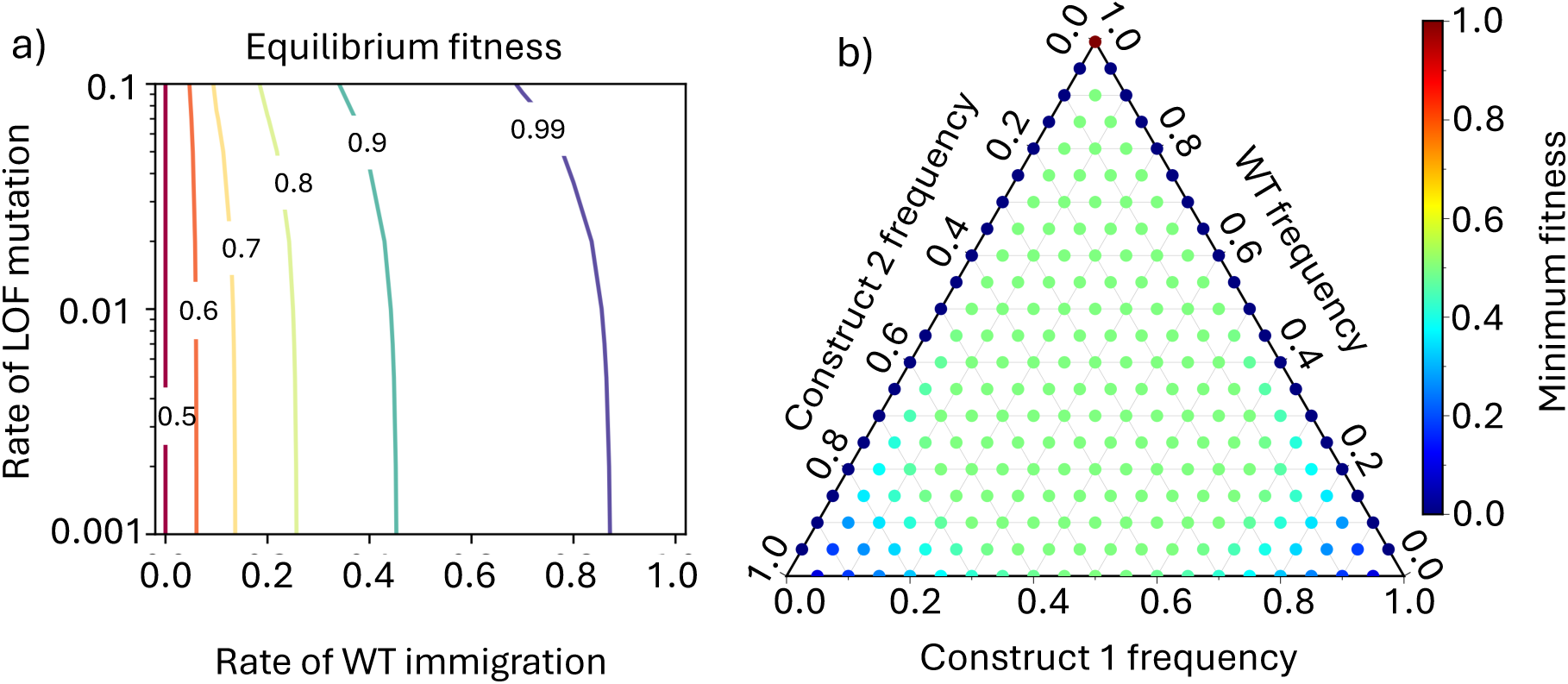
**a)** Equilibrium fitness of a population under a balanced lethal system as a function of the rate of wild-type immigration and the rate of homing-associated loss-of-function (LOF) mutation. Contour lines indicate equilibrium fitness value, where 0.5 corresponds to the maximum 50% genetic load (no immigration, no loss-of-function mutation) and 1.0 corresponds to no genetic load. **b)** Minimum fitness experienced during the trajectory to equilibrium from different starting combinations of wild-type and two homing construct allele frequencies. Each point represents a simulation initiated at a different starting allele composition. Colour indicates the minimum fitness (maximum genetic load) experienced at any point during the trajectory. All simulations assume 100% homing, no loss-of-function mutation and no migration.

Before the balanced lethal equilibrium is reached, transient dynamics during establishment can also depart from the idealised case. When allele frequencies of the two constructs are unequal, the load can temporarily exceed 50% before returning to 50:50 equilibrium frequency, as a greater proportion of homozygotes for the more common construct are formed (Figure 4b). Similar transient overshoots arise when a resistant allele is released in place of a homing one, and can be more pronounced because the resistant allele lags behind the homing construct during spread (Supp. Figure 2). In the extreme case where one of the constructs is lost from the population entirely, for example due to stochastic effects in a small population, the remaining homing construct can spread to fixation, since no complementing allele remains to maintain the balanced lethal system. This can lead to substantially increased load that may be sufficient for population elimination (Figure 4b).

### Localisable homing designs

The designs explored above are expected to spread from rare to their equilibrium frequency and therefore have the potential to spread across a species’ entire range wherever there is gene flow. Where isolation is required, this can be achieved using a split-drive design, in which the Cas9 is removed from both balanced lethal constructs and instead placed at a separate neutral autosomal locus (Supp. Figure 5), analogous to previous split homing drives for population replacement (Noble et al., 2019). The balanced lethal alleles can then home only in the presence of Cas9. Because Cas9 is inherited in a Mendelian fashion and does not itself spread, it cannot establish in neighbouring wild-type populations where it arrives at low frequency through migration, confining the balanced lethal system to the target population (Figure 5).

**Figure 5.**
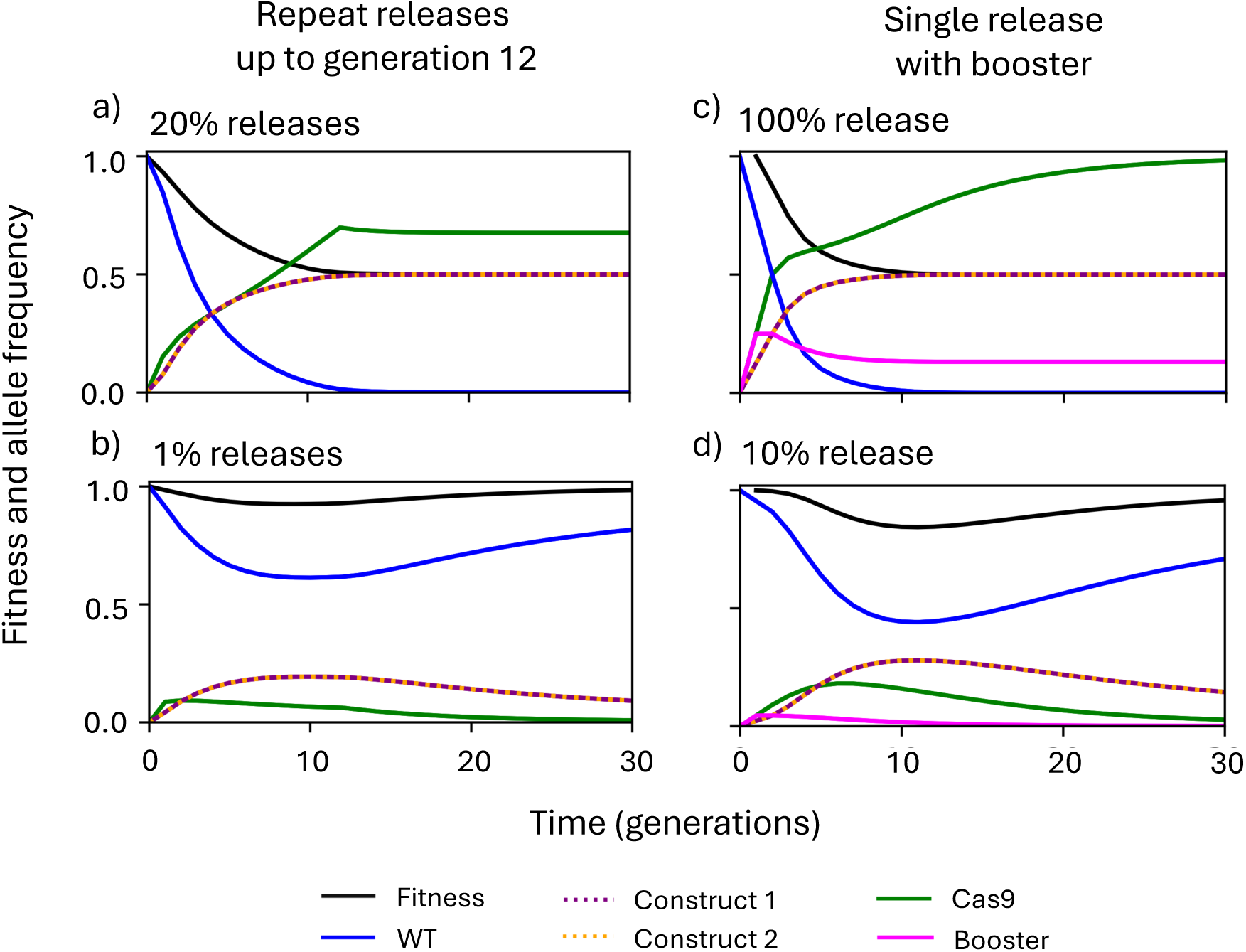
Establishment of a localised balanced lethal system using a split-drive design. In all panels, the Cas9 editor is located at a separate autosomal locus from the balanced lethal alleles. Lines show population mean fitness (black), wild-type allele frequency (blue solid), construct 1 and 2 (purple and orange dashed respectively, overlayed), Cas9 editor (green solid), and booster (pink solid). **a)** Repeated releases of trans-heterozygous males (homozygous for the editor) at 20% of the male population each generation for 12 generations. The balanced lethal system is established: both construct alleles reach 50% frequency, the wild-type is eliminated, and a sustained 50% load is achieved. **b)** As in (a), but with 1% releases. The release size is insufficient to establish the balanced lethal system and the wild-type allele is not eliminated. **c)** A single release at 100% of the male population, with the editor paired with a booster construct containing a gRNA at a third locus (magenta solid) that causes the Cas9 to home. The booster temporarily increases editor frequency, enabling elimination of the wild-type and establishment of the balanced lethal from a single release. **d)** As in (c), but with a 10% single release, where the release size is insufficient for establishment.

As with the autonomous designs, the balanced lethal system is self-maintaining once the wild-type allele is eliminated, since the only viable adults are trans-heterozygotes. Establishment therefore requires that the wild-type is fully eliminated before the Cas9 is lost from the population. Repeated releases of trans-heterozygous males homozygous for Cas9 progressively push the balanced lethal alleles to higher frequency, and if releases are sufficiently large and sustained, the wild-type is eliminated and the system establishes (Figure 5a). Smaller releases fail to establish (Figure 5b), and it is for this reason low-level emigration of the constructs into neighbouring wild-type populations would not result in establishment there, confining the system to the release site. To reduce the release effort required for establishment, the Cas9 can be paired with a booster construct, a gRNA at a third locus that causes the Cas9 to home (Supp. Figure 5), temporarily increasing its frequency and enabling establishment from a single release (Figure 5c). As with the non-boosted design, releases below a threshold size fail to establish (Figure 5d). An alternative route to localisation would be to pair the out-of-locus Cas9 with a gRNA targeting a geographically restricted sequence, allowing it to self-amplify in the target population while being unable to spread where that sequence is absent, analogous to double-drive designs proposed for population suppression (Willis and Burt, 2021).

In both boosted and non-boosted designs, the Cas9 at its own locus is assumed to be selectively neutral in the absence of homing. However, while wild-type alleles remain at the balanced lethal locus, homing driven by the Cas9 converts them to construct alleles, increasing the frequency of lethal homozygotes among offspring. This indirectly reduces the effective fitness of Cas9-carrying individuals, since their offspring are more likely to inherit a construct allele and die as homozygotes. Once the wild-type at the balanced lethal locus is eliminated, homing ceases and the Cas9 becomes selectively neutral. In the presence of immigration, however, wild-type alleles are continually reintroduced at the balanced lethal locus, continually reimposing this indirect fitness cost on the Cas9 and causing it to decline (Supp. Figure 4). Suppression is therefore likely to be temporary in connected populations. This mechanism provides a route to deliberate reversal: releasing wild-type individuals would reimpose the fitness cost on the Cas9, causing it to decline and the population to recover.

## 3. DISCUSSION

### Summary

Our modelling shows that two complementing homing constructs released at low frequency can establish a balanced lethal system, eliminating the wild-type allele and imposing a sustained 50% load through Mendelian segregation. The population-level consequences depend on the strength and timing of density-dependent mortality: the same load can suppress populations, increase them, or dampen oscillatory dynamics, with the direction and magnitude of the effect determined by Rm and B. Additional systems at independent loci scale the effect in stepwise increments, and a split-drive variant confines establishment to the target population through a release threshold. Because the load arises from the knockout phenotype of two characterised alleles rather than from the field fitness of a partially disrupted gene, its magnitude is more predictable than in existing partial-suppression designs. Together, these properties extend load-inducing gene drives beyond pest elimination, with applications in conservation and management of beneficial and endangered populations.

### Resistance and robustness

Resistance alleles are a general concern for all gene drives (Unckless et al., 2017, Champer et al., 2017, Kyrou et al., 2018). In this system, loss-of-function resistance alleles, those that disrupt the target gene without restoring wild-type function, are benign, provided they maintain the knockout phenotype and participate in the balanced lethal system (Figure 4a). Functional resistance alleles that restore wild-type function whilst preventing homing would, by contrast, be expected to spread and prevent establishment. Generating such alleles would require precise in-frame repair at the target site, which may be minimised by targeting conserved sequences and by multiplexing gRNAs at each target site, so that resistance must arise at all sites simultaneously to prevent homing (Kyrou et al., 2018, Fuchs et al., 2021, Yang et al., 2021, Oberhofer et al., 2018, Champer et al., 2018).

A second route to resistance is recombination between the two insertion sites in trans-heterozygotes, which could generate a haplotype carrying both recoded target sequences: a functional allele resistant to homing by either construct. The reciprocal product, carrying both disruptions, would be lethal and therefore eliminated. The rate of such events depends on the physical distance between the two target exons; selecting a gene in which they are in close proximity would both minimise this risk and ensure that the recoded target site is co-transferred with the construct during homing. Shared sequence homology between the two constructs (for example, Cas9 cassettes) could also facilitate non-allelic recombination between them. This could be avoided by using distinct codon-optimised sequences in each construct, or by adopting the single homing-construct design released alongside a non-driving complementary allele, which minimises shared sequence between the two alleles.

A further consideration arising from using two complementing constructs at separate sites is the length of the gene conversion tract during homing. The design requires that each construct co-transfers a recoded version of the other construct’s target site, so that the homed chromosome is resistant to cleavage by both gRNAs. If the gene conversion tract is shorter than the distance between the two sites, this recoded site would not be co-transferred, allowing cross-homing in trans-heterozygotes and the formation of haplotypes carrying both disruptions. Such haplotypes would not be a source of resistance, since they are lethal and would be selected against, but their formation represents a route to reduced establishment efficiency.

Beyond resistance, our modelling assumed complete homing efficiency and no heterozygous fitness costs of the constructs. Imperfect homing would slow establishment but would not affect the equilibrium, since the balanced lethal system depends on the knockout phenotype rather than on the drive machinery. Heterozygous fitness costs would impede spread, and if sufficiently large could prevent elimination of the wild-type. Even where the wild-type persists, however, the reduced fitness of construct-carrying heterozygotes would itself impose a load and could still reduce population size. The population-level consequence would then depend on the timing of these heterozygous costs relative to density-dependent mortality, in the same way as for the lethality of homozygotes. Because the two need not act at the same life stage, lethality and heterozygous costs could be imposed before and after density dependence in any combination, and the net effect on population size would reflect their interaction. The sensitivity of the system to resistant allele formation, homing efficiency and fitness parameters, and any interaction between them, warrants further modelling.

### Alternative modes of intragenic complementation

In our results, we illustrated the approach with one architecture, a gene producing two essential isoforms through alternative splicing, but intragenic complementation can arise through several mechanisms. It has been observed in (i) genes that encode proteins with multiple functional domains, where mutations affecting different domains allow trans-heterozygotes to maintain function from the intact domains of each allele (e.g. *rudimentary* in *Drosophila (Carlson, 1971)*, *let-2* in *C. elegans (Sibley et al., 1993))*; (ii) genes producing different isoforms through alternative splicing or alternative promoters, where mutations in different isoforms allow trans-heterozygotes to express the full complement (e.g. *Sex lethal* in *Drosophila (Salz et al., 1987))*; and (iii) genes that undergo trans-splicing, where intact exons transcribed from different chromosomes combine into functional transcripts (e.g. *longitudinals lacking* in *Drosophila (Horiuchi et al., 2003))*.

The engineering approach for constructing a balanced lethal system differs by mechanism. For isoform-based (ii) and trans-splicing-based (iii) complementation, premature stop codons placed in exons specific to different isoforms or splice units could create the complementing pair, with the homing cassette inserted immediately downstream. For domain-based complementation (i), missense mutations disrupting specific functional domains while preserving protein structure would be required, since premature stop codons would eliminate all downstream domains; the homing machinery could then be placed in a nearby intron, with the gRNA targeting the wild-type sequence at the mutation site so that both mutation and cassette are copied together. Across all designs, the wild-type target sequence must be absent from both complementing alleles to prevent cross-homing in trans-heterozygotes, Cas9 should be expressed from a germline promoter to avoid creating homozygotes in the soma, and any insertions must preserve the splice patterns of the unaffected isoform. The target gene must also tolerate a reduction in expression in trans-heterozygotes, since each allele contributes only half of the gene’s functional output, making haplo-insufficient genes unsuitable targets.

If a 50% load is stronger than required, genes amenable to three or more complementing alleles could in principle be used to engineer three or more constructs at a single locus. With *n* alleles at equal frequency, the genetic load from homozygote lethality is 1/*n*, giving approximately 33% load with three alleles and 25% with four. This would require all pairwise combinations to complement, which may be achievable for genes with three or more independent functional domains or isoforms. Some genes are known to support multiple complementarity groups. In *Drosophila melanogaster*, *rudimentary (Carlson, 1971)* and the *Broad complex (Kiss et al., 1988)* each exhibit three complementing groups, while *lola* has at least 80 different protein isoforms, with a range of different constant and variable exons exhibiting a complex complementarity patterns *(Horiuchi et al., 2003)*.

### Alternative molecular designs

We focused our results on homing-based drive mechanisms, but the same balanced lethal equilibrium can be reached using other drive mechanisms, for example CRISPR-based toxin antidote (TA) drive. If capable of eliminating the wild-type, these would differ only in the transient dynamics during spread; the equilibrium state and its consequences for population dynamics would be the same. In one approach, each of two complementing knockout alleles is paired with a cleave-and-rescue element: a Cas9 and gRNA that create dominant lethal edits at an unlinked essential gene (cleave), linked to a recoded copy of that gene that rescues the lethality (rescue). Both constructs must target the same essential gene: if they target different genes, offspring of released trans-heterozygous males would inherit both toxins and none would survive. With both constructs targeting the same gene, each complementing allele would spread from rare through TA drive, eliminating the wild-type and establishing the balanced lethal system. Both the homing and TA designs described so far require intragenic complementation at a single locus, constraining the choice of target gene. It may be possible to engineer this at two loci, more closely resembling mechanisms found in nature and relaxing the requirement for the complementing alleles to be different functional variants of the same gene, though at the expense of a more complex design.

As with the homing approach, a single active-construct design is possible, closely related to the protected dominant-negative editor (PDNE) (Willis and Burt, 2025a), and potentially simpler to engineer. In the PDNE, a construct creates dominant lethal edits and carries a cis-acting rescue element to achieve drive. The construct itself is recessive lethal, creating a fitness cost that precisely balances the drive and resulting in a selectively neutral construct that persists at its release frequency. If the rescue instead acts in both cis and trans, using for example RNAi, construct carriers would be fully protected, the drive would be strengthened, and the construct would increase in frequency. Because the construct itself is a recessive lethal, once the wild-type is eliminated only construct/edit heterozygotes survive (through trans rescue), while homozygotes for either the construct or the edit allele are lethal, establishing a balanced lethal system.

### Comparison with other approaches to partial suppression

Several alternative strategies could in principle be used to achieve partial suppression. Non-driving approaches such as the sterile insect technique, RIDL or non-driving editors impose load dependent on release rate, but require relatively large and continuous releases to maintain significant suppression (Alphey et al., 2010, Dyck et al., 2021, Willis and Burt, 2025b). Selectively neutral designs such as the Y-linked editor, male-drive female-sterile and PDNE allow the level of load to be tuned through release size, and once established, do not require repeat releases, since they are self-sustaining in the absence of costs. Unlike the balanced lethal design, however, they cannot establish by self-amplifying from rare. Where this spread is undesirable, our split-homing variant provides confinement through a threshold release for wild-type elimination. A precise comparison of establishment efficiency between these self-sustaining designs and the split balanced lethal system is left to future work.

The closest relatives of the design proposed here are the UDmel system of Akbari et al. (2013) and the 1-locus 2-drive TARE design of Champer et al. (2020a), both of which pair two reciprocal toxin-antidote constructs at a single locus to produce 50% balanced lethality. UDmel comprises two maternally-expressed toxin-antidote pairs, with each antidote linked to the non-cognate toxin, such that viable transgenic offspring must inherit both constructs. The TARE design uses two TA constructs at the same locus, each targeting a different haplo-sufficient gene and providing rescue for the other’s target, with the reciprocal cleave-and-rescue itself generating the balanced lethality. Unlike the designs presented here, where the drive machinery and the balanced lethal phenotype are mechanistically independent, the UDmel and TARE designs couple drive and balanced lethality through a single reciprocal-rescue architecture. Further modelling would be required to compare each of these strategies in terms of their release requirements, spread dynamics, localisation, resilience to resistance and performance in the face of wild-type immigration.

### Conclusions

We have shown that a balanced lethal system can be engineered through gene drive to impose a sustained 50% genetic load that, depending on a population’s density regulation, can suppress, enhance, or stabilise it. Realising these applications in any particular species will require empirical characterisation of its density regulation and the timing of density-dependent mortality. While the genetic load imposed by the balanced lethal system is predictable by design, its population-level consequence is not, and methods for estimating density dependence from field data (Hassell et al., 1976, Bellows, 1981, Turchin, 2003) will be important tools for strategy choice and deployment planning. This requirement is especially acute for boosting applications, which require overcompensating density regulation and lethality acting before density-dependent mortality, and dampening oscillating populations such as locusts, where release timing relative to natural cycles becomes a potential design variable. Further investigation into the regulatory dynamics of natural populations, alongside experimental construction of the designs themselves, will therefore be essential steps towards deployment.

## 4. METHODS

To simulate the dynamics of a balanced lethal system established through gene drive, we developed a series of deterministic models simulating a single, panmictic population of infinite size with discrete, non-overlapping generations, based on the framework described in Burt and Deredec (2018). The baseline model has a single autosomal locus with three alleles: a wild-type allele (+) and two construct alleles (A and a), each inserted into a different exon of the same essential gene. Males and females are modelled separately, giving six genotypes per sex. Each construct disrupts one essential protein isoform while preserving the other, so homozygotes for either construct (A/A or a/a) are lethal, while trans-heterozygotes (A/a) are viable through intragenic complementation. Wild-type heterozygotes (A/+ and a/+) and wild-type homozygotes (+/+) are also viable and assumed to have full fitness.

Each construct contains a Cas9 and a guide RNA (gRNA) targeting the wild-type sequence at its own insertion site, enabling homing via homology-directed repair. Each construct also carries a recoded version of the target sequence at the other exon, preventing homing between the two constructs in trans-heterozygotes. In genotypes containing at least one construct allele and one wild-type allele at the same locus, homing converts the wild-type allele to the construct allele with probability *c*. No difference in homing rate was assumed between individuals carrying one or two copies of the construct. The fitness effects of the constructs are modelled by parameters *s* and ℎ, where *s* is the fitness cost in homozygotes and *h* is the dominance coefficient giving the proportion of homozygous cost experienced by heterozygotes. For the default balanced lethal parameterisation, *s* = 1 and ℎ = 0 for both construct alleles, with fitness of trans-heterozygotes set to 1. In all baseline results, homing efficiency was set to *c* = 1 and no NHEJ resistant products were formed.

To model loss-of-function (LOF) mutations associated with the homing machinery, each construct component (Cas9 or gRNA) has a probability *d* per generation of becoming non-functional. Degenerate alleles retain their knockout phenotype and complementarity, and are therefore functionally equivalent to their intact counterparts in the balanced lethal system, but can no longer convert wild-type alleles through homing. To model immigration, wild-type males and females enter the population each generation at a rate *m*, expressed as a proportion of the current population.

To model a localisable balanced lethal system, the baseline model was extended to include a second autosomal locus carrying two alleles, the wild-type and a construct containing the Cas9. The balanced lethal alleles at the first locus carry only the gRNAs, and can therefore home only in the presence of the Cas9 at the second locus. The two loci are assumed to be unlinked. To model the Cas9 paired with a booster construct, a third autosomal locus was included, carrying two alleles, the wild-type and a booster construct containing a gRNA that targets the wild-type allele at the Cas9 locus. In the presence of both the booster and a wild-type allele at the Cas9 locus, the Cas9 homes with probability *c*, converting the wild-type to a Cas9 allele. The booster was assumed to be unlinked from both the Cas9 and balanced lethal loci, and non-autonomous (unable to function in the absence of the Cas9). To model multiple independent balanced lethal systems for stepwise population suppression, the model was extended to include a second independent autosomal balanced lethal locus, unlinked from the first. Each locus operates as described above, with its own pair of complementing construct alleles and independent homing dynamics.

To evaluate the impact on population size, we coupled the genetics to a population dynamics model tracking the number of males and females over time, through egg, hatchling, pupal and adult stages. In each generation, females produce *f* eggs, which are fertilised by males, assuming male numbers do not limit fertilisation success and that mating is random. Density-dependent mortality is applied at the hatchling-to-pupal transition according to a generalised Beverton-Holt model, in which survival from larvae to pupal stage is:

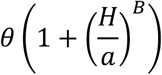

where *θ* is the density-independent probability of surviving the larval stage, *B* determines the strength of competition, *a* sets the scale at which density-dependent mortality occurs and *H* is the total number of larvae in the population. Since all results are reported as population size relative to the starting population, the value of *a* did not matter. The intrinsic rate of increase is 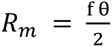. All releases were of adult males, either trans-heterozygous for the balanced lethal alleles alone, or additionally homozygous for the Cas9 and/or booster. Allele frequencies were reported at the hatchling stage, calculated in males and females independently and then averaged. All simulations were performed in Julia within the Jupyter notebook environment.

## ACKNOWLEDGEMENTS

We would like to thank John Connolly, Silke Fuchs and Alekos Simoni for their valuable feedback. This work was supported by the Gates Foundation.

## Notes

### Competing Interest Statement

The authors have declared no competing interest.

