## Supplementary Material for "Engineered balanced lethal systems for partial suppression or enhancement of wild populations"

### Supplementary Figures

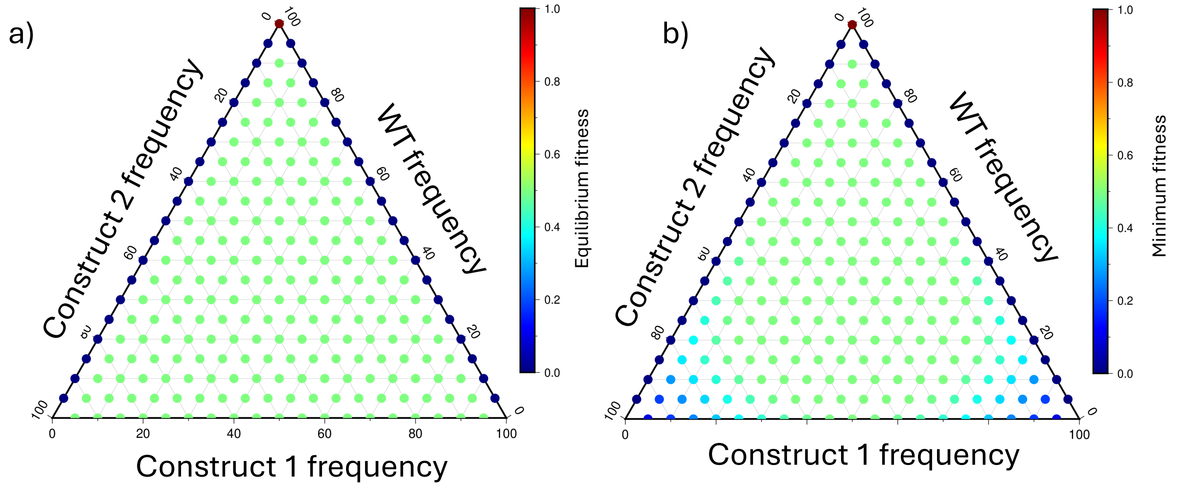


***Supp. Figure 1 - a)*** *The equilibrium allele frequency following different start state allele frequencies.* ***b)*** *The minimum allele frequency following different start state allele frequencies.*

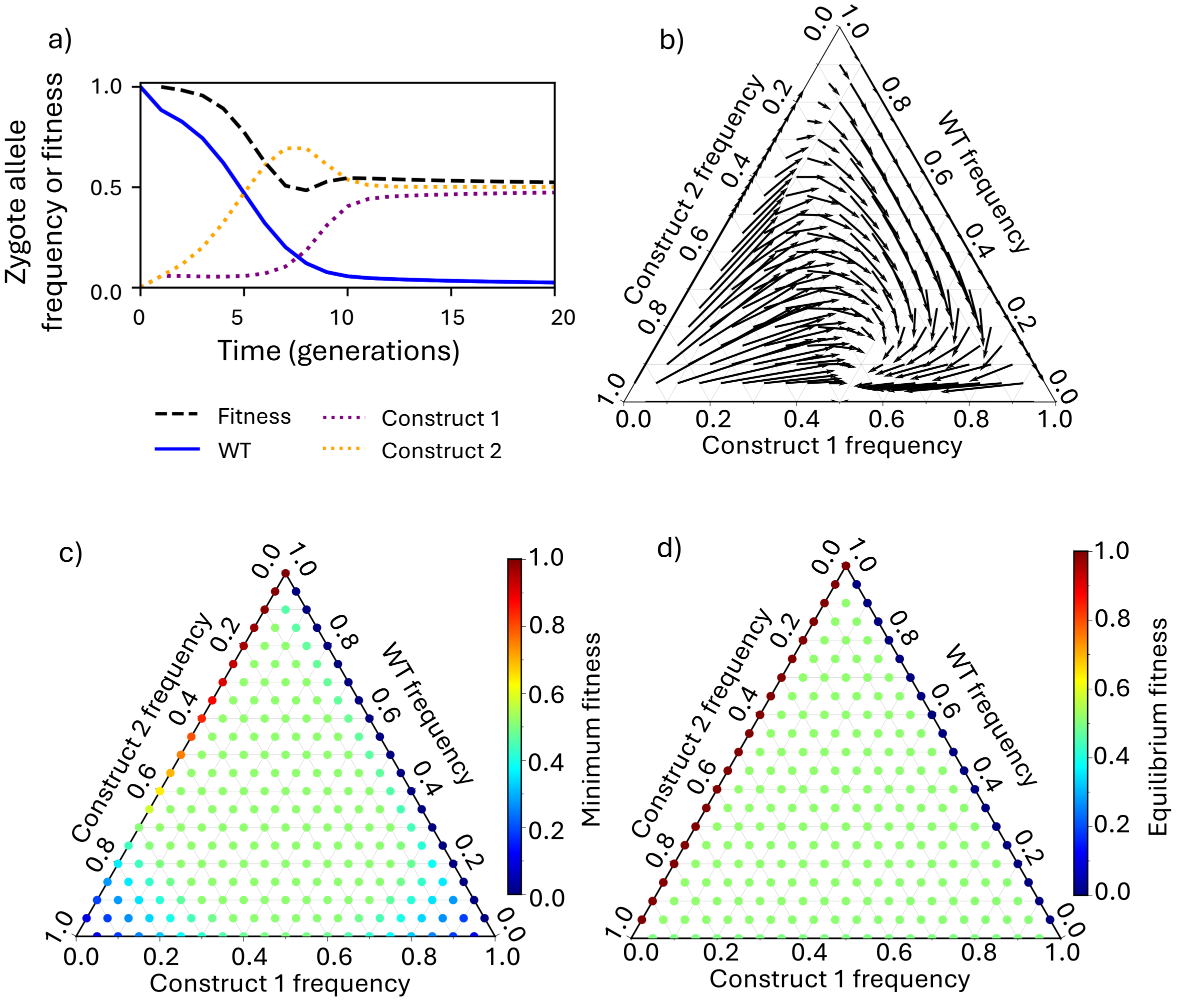


***Supp. Figure 2** – Homing balanced lethal system where one allele homes (orange) and the other doesn't (purple). All other labels are as in Figure 1 and supplementary figure 1.*

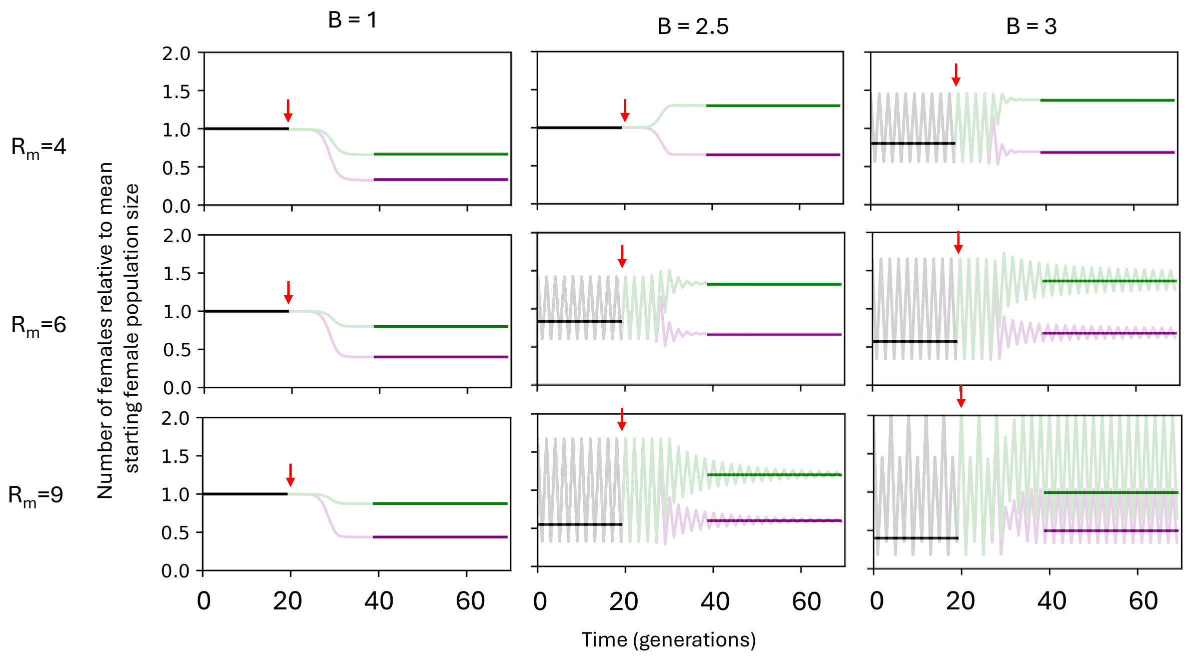


***Supp. Figure 3 –** As Figure 2, showing additionally the harmonic mean at equilibrium before (black) and after intervention (green and purple). Note that the pre-release harmonic mean is below 1 than B > 1 because it is calculated from a fluctuating population: the harmonic mean is always less than or equal to the arithmetic mean when census size varies over time.*

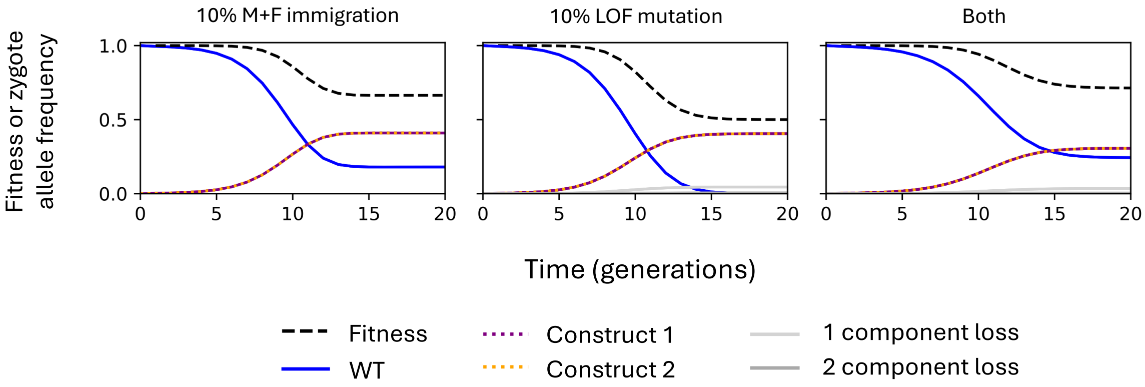


***Supp. Figure 4*** *– Time course of a release of a balanced lethal system in the presence of wild-type immigration each generation and loss of function mutations. Releases are of two homing gene drives released in heterozygous males carrying one copy of each type at generation zero at 1% of the initial male population size. Lines show allele frequencies of the wild-type (blue), construct 1 (purple) and construct 2 (orange), the frequency of construct alleles that have lost one (light grey) or both (dark grey) homing components through loss-of-function mutation, and mean population fitness (dashed black). Loss-of-function alleles retain the knockout phenotype and continue to participate in the balanced lethal system but can no longer convert wild-type alleles by homing.*

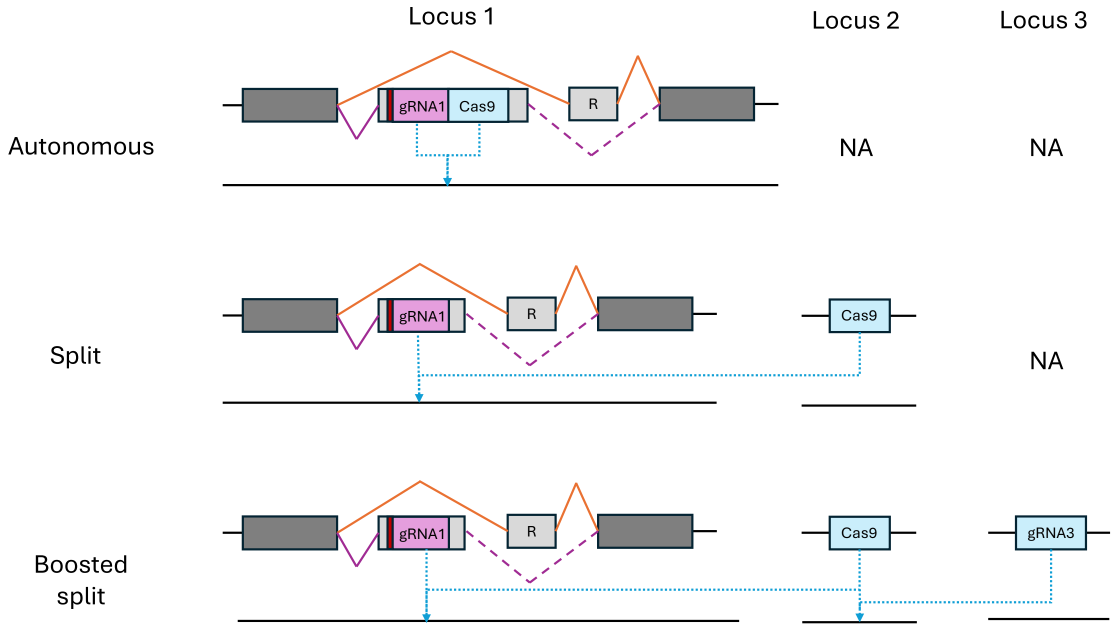


***Supp. Figure 5** – Schematic of the three balanced lethal designs. Each row shows the construct alleles present at each locus for the autonomous (top), split (middle), and boosted split (bottom) designs. Locus 1 carries one of the balanced lethal constructs; in the autonomous design this also contains Cas9 (analogous to Figure 1a). The recoded target site (R) prevents homing between the two complementing constructs in trans-heterozygotes. Orange and purple lines indicate the two alternative splice patterns; dashed lines indicate disruption of splicing by construct insertion. Blue dotted arrows indicate CRISPR-based cleavage and homing. In the split and boosted split designs, Cas9 is located at a separate neutral locus (Locus 2). In the boosted split design, a non-autonomous booster construct inserted into Locus 3 contains a gRNA that targets the WT allele at locus 2, allowing Cas9 to home into the wild-type allele. gRNA3 requires Cas9 to function, preventing it from acting on wild-type alleles in the absence of Cas9. For clarity, Locus 1 shows only Allele 1, which disrupts isoform 1 and carries gRNA1. In the full system, a complementing Allele 2, disrupting isoform 2, is also present at Locus 1; together the two alleles form the balanced lethal system. For simplicity, the splicing pattern of the WT allele is omitted, but is analogous to that shown in Figure 1a.*
